# Time-course Profiling of Bovine Herpesvirus Type 1 and Host Cell Transcriptomes using Multiplatform Sequencing

**DOI:** 10.1101/2020.05.25.114843

**Authors:** Norbert Moldován, Zoltán Maróti, Gábor Torma, Gábor Gulyás, Ákos Hornyák, Zoltán Zádori, Victoria A. Jefferson, Zsolt Csabai, Miklós Boldogkői, Tibor Kalmár, Dóra Tombácz, Florencia Meyer, Zsolt Boldogkői

**Affiliations:** Department of Medical Biology, Faculty of Medicine, University of Szeged, Somogyi B. u. 4., 6720 Szeged, Hungary; Department of Pediatrics, Faculty of Medicine, University of Szeged, Somogyi B. u. 4., Szeged, H-6720, Hungary; Institute for Veterinary Medical Research, Centre for Agricultural Research, Hungarian Academy of Sciences, Hungária krt. 21, H-1143 Budapest, Hungary; Dept. of Biochemistry & Molecular Biology, Entomology & Plant Pathology, Mississippi State University, 408 Dorman Hall - Box 9655, 32 Creelman St., Starkville MS 39762, USA

**Author notes:** The first two authors contributed equally to this work. NM ZM GT GG ÁH ZZ VJ ZC MB TK DT FM ZB.

**Keywords:** herpesviruses, bovine herpesvirus type 1, long-read sequencing, direct RNA sequencing, Oxford Nanopore Technology, MinION, Illumina, LoopSeq, transcript isoforms, MDBK cells

## Abstract

Long-read sequencing (LRS) has become a standard approach for transcriptome analysis in recent years. This technology is also used for the identification and annotation of genes of various organisms, including viruses. Bovine herpesvirus type 1 (BoHV-1) is an important pathogen of cattle worldwide. However, the transcriptome of this virus is still largely unannotated. This study reports the profiling of the dynamic lytic transcriptome of BoHV-1 using two long-read sequencing (LRS) techniques, the Oxford Nanopore Technology (ONT) MinION, and the Illumina LoopSeq synthetic LRS methods, using multiple library preparation protocols. In this work, we annotated viral mRNAs and non-coding transcripts, and a large number of transcript isoforms, including transcription start and end sites, as well as splice variants of BoHV-1. Very long polycistronic and complex viral transcripts were also detected. Our analysis demonstrated an extremely complex pattern of transcriptional overlaps formed by transcriptional read-throughs or overlapping the 5’-untranslated regions of divergently-oriented transcripts. The impact of the viral infection on the host cell transcriptome was also assessed. Our results demonstrate that genes associated with antiviral response as well as viral transcription and translation are upregulated.

## INTRODUCTION

Bovine herpesvirus type 1 (BoHV-1) belongs to the subfamily Alphaherpesvirinae of the family Herpesviridae. BoHV-1 largely infects cattle and related ruminants (Thiry et al., 2006). The viral infection leads to immune suppression and conjunctivitis, which allows bacteria to invade the lower respiratory tract leading to pneumonia. BoHV-1 is one of the viruses that triggers the disease commonly known as bovine respiratory disease (BRD), which costs the cattle industry several billion dollars annually worldwide (van Oirschot, 1995). BoHV-1 acute infection induces apoptosis in CD4^+^ T cells (Winkler et al., 1999), impairs antigen processing and CD8^+^ T cell recognition of infected cells (Hinkley et al., 1998) and dampens the interferon type I response through multiple mechanisms (Henderson et al., 2005). Similarly to other alphaherpesviruses, such as herpes simplex virus type 1 (HSV-1), or pseudorabies virus (PRV), BoHV-1 also enters a latent state especially in the trigeminal ganglia of the host following primary infection (Jones, 1998), from this state the virus can be reactivated by various types of stress, and re-establish an acute infection (Nataraj et al., 1997). BoHV-1 is a large (∼136 kbp) double-stranded DNA virus. The viral genome has been sequenced and structurally annotated by an international effort more than twenty years ago, using viral DNA from several strains/subtypes (Khattar et al., 1995; Schwyzer et al., 1996; Vlček et al., 1995). Full sequencing of individual strains has since occurred for BoHV-1.1 (d’Offay et al., 2013), BoHV-1.2b (d’Offay et al., 2016),[and several vaccine and field isolates (Fulton et al., 2013). Yet, to date the original genomic annotation of 69 open reading frames (ORFs) has not been revised. Several new ORFs are now recognized to exist in and around the latency-related (LR) gene (Meyer et al., 2007a), and recently a new ORF was identified using experimental proteomic data (Meyer et al., 2007b). Proteogenomic data has also revealed at least 92 unannotated peptides coded by the BoHV-1.1 genome, 21 of which are surrounded by potential ORFs identified *in silico* (Jefferson et al., 2018). The herpesvirus genes can be classified to immediate-early (IE), early (E), early-late (L1) and late (L2) genes depending on the kinetics they are expressed throughout the viral replication cycle (Harkness et al., 2014). The IE genes are typically the transactivators of many other viral genes, the E genes encode the enzymes needed for the DNA synthesis, whereas the L genes specify the structural elements of the virions. ORFs of the BoHV-1 genes are mainly annotated *in silico*, whereas the size and the possible splicing patterns of many transcripts was detected using Northern blot and S1 nuclease mapping (Schwyzer et al., 1993, 1994; Vlček et al., 1995; Wirth et al., 1992). Until now, the only genome-wide transcriptome assay focused on the detection of microRNAs of BoHV-1 (Glazov et al., 2010).

Next-generation sequencing (NGS) [short-read sequencing (SRS)] technology has revolutionized transcriptome research due to its capacity to sequence a large number of nucleic acid fragments simultaneously, at a relatively low cost. In the last couple of years, third-generation sequencing (TGS) [long-read sequencing (LRS)] has become an alternative approach that is able to circumvent the limitations of SRS, including its inability to read full-length RNA molecules that are necessary, among others, for the identification of transcript isoforms and for distinguishing between overlapping transcripts. Recently, TGS has been widely applied for the transcriptome analysis of a variety of organisms (Boldogkői et al., 2019; Byrne et al., 2017; Chen et al., 2017; Cheng et al., 2017; Jiang et al., 2019; Li et al., 2018; Moldován et al., 2017, 2018a, 2018b; Nudelman et al., 2018; Tombácz et al., 2018a; Zhang et al., 2018; Zhao et al., 2019), including herpesviruses (Balázs et al., 2017; Depledge et al., 2019; Moldován et al., 2018a; O’Grady et al., 2016; Tombácz et al., 2015, 2016, 2017, 2018b). These approaches have revealed a much more complex transcriptional landscape of the examined viruses than previously thought.

In this study, we carried out a time-lapse analysis of BoHV-1 and host cell transcriptomes using two LRS techniques, ONT MinION and Loop Genomics LoopSeq™ synthetic long-read sequencing on the Illumina platform.

## RESULTS

### Time-lapse transcriptome analysis using long-read sequencing

In this work, we carried out a time-course analysis of the BoHV-1 transcriptome and of the impact of the viral infection on host cell gene expression. In order to capture the most complete transcriptome dataset, we applied various experimental conditions, including two different cell types for virus propagation, different multiplicities of infection, various library preparation methods and different sequencing platforms. Transcriptome analysis was performed using two LRS technologies: ONT MinION sequencing and LoopSeq synthetic long-read sequencing on the Illumina platform. We applied both cDNA and direct RNA sequencing, as well as oligo(dT) and random oligonucleotide-primed reverse transcription for the ONT platform. We also used amplified and non-amplified cDNA-based techniques. ONT sequencing produced altogether 3,387,951 sequencing reads mapping to the BoHV-1 genome. LoopSeq analysis generated 36,627,542 sequencing reads, which were assembled into 27,098 contigs, representing full-length reads. Of these, 12,618 mapped to the viral genome. The detailed summary statistics is presented in **Supplementary information**, Error! Reference source not found.. In previous publications, we (Moldován et al., 2017b, 2018b) and others (Workman et al., 2018) have reported that the direct RNA sequencing (dRNA-Seq) method is not suitable for capturing complete transcripts, since short (15-30bps) sequences from the 5’ region and in many cases also the poly(A)-tails were missing from the reads. The lack of 5’-ends is the result of the release of the RNA strand from the ratcheting protein leading to the rapid transition of RNAs through the pore. On the other hand, polyA-tails are probably missing due to the presence of a DNA adapter ligated to the 3’-end during library preparation. This adapter muddles the signal near the 3’-ends of the RNA molecule, resulting in erroneous base calling. Another drawback of native RNA sequencing is its relatively low throughput compared to cDNA sequencing. On the other hand, dRNA-Seq is free of artefacts produced by RT, second strand synthesis, and PCR. We used random oligonucleotide-primed cDNA synthesis for TSSs and splice site validation. Three biological replicates were prepared for each time-point in the dcDNA sequencing experiment. Altogether, 29 reactions were run using 5 different techniques for providing independent reads. The RT, the second strand synthesis and the PCR often result in nonspecific binding of oligod(T) or PCR primers and in template switching. It has been demonstrated that oligo(dT) primers can occasionally hybridize to A-rich regions of the mRNA or to the first cDNA strand and thereby producing false transcription end sites (TESs) or truncated 5’-ends (Sessegolo et al., 2019). Such products were eliminated from further analysis by the LoRTIA software suite, by checking the presence of 5’ sequencing adapters on the read, and homopolymer A-s in the near vicinity of read ends on the genome. The LoRTIA toolkit detected a total of 2,870 putative transcription start sites (TSSs), 475 putative TESs, and 645 putative introns in the MinION sequencing data **(Table S2)**. TSS and TES were accepted if the LoRTIA pipeline detected them in at least three independent samples. Applying this filter resulted in 823 TSSs and 135 TESs, which were used for transcript annotation. Altogether, 25 introns were identified using our filtering criteria, among which 23 carried canonical GT/AG and 2 GC/AG splice junction sequences **(Table).**

The selected TSSs, TESs and introns, used for transcript annotation, yielded a total of 1,381 putative transcripts. To reduce the number of possibly false isoforms, we excluded putative transcripts detected in less than three samples, which resulted in a total of 1,025 transcripts **(Table).**

Our investigations revealed that the entire BoHV-1 genome is transcriptionally active (including the genomic junction) with the exception of a 336-nt region between the TESs of CIRC and UL54, as well as a 1,638-nt genomic segment between the TSS of bICP4 and the TES of ORIS-RNA1.

### Sequencing of BoHV-1 genome

In this study, we used the Cooper strain of BoHV-1 maintained in two different laboratories (in Hungary and in Mississippi, USA). We carried out native DNA sequencing in order to monitor the potential effect of mutations accumulated throughout propagation on the gene expression, and to exclude that the annotated splice sites derived from genomic deletions.

#### The BoHV-1/MS genome

The sequencing of the BoHV-1/MS sample resulted in a total of 46,922 reads which were assembled into a 153,076-nt long contig. This was then aligned to the reference genome, and contig ends were trimmed, resulting in a final genome length of 137,652 nt. To assess the assembly, we mapped the sequencing reads to the assembly. 11,741 reads mapped to the assembled viral genome with an average read length of 4,955 nt (σ=7,622) and an average coverage of 423.

#### The BoHV-1/HU genome

The sequencing of the BoHV-1/HU sample yielded a total of 39,205 reads that were assembled and polished into a 137,550-nt long contig. We assessed the assembly by mapping the reads to the assembled contig. 17,138 reads mapped to this assembly with an average read length of 8,379 nt (σ= 5,372) and with an average coverage of 1,044.

Based on these alignments, we concluded that differences found between the reference and the two assembled viral genomes do not affect the exon composition of the transcripts.

### Novel putative protein-coding genes: embedded genes

In principle, all of the transcripts should be considered as new because conventional techniques were unable to detect full-length transcripts and therefore match identified TSS and TES sequences. However, we only consider the transcripts containing 5’-truncated in-frame ORFs as novel putative mRNAs. These transcripts are produced by promoters located within larger canonical ORFs, or theoretically, in those cases where no promoters were identified by polymerase slippage, a mechanism which – among herpesviruses - has only been described in KSHV (Atkins et al., 2016). LRS techniques have proven to be very useful for the detection of the hidden complexity in embedded ORF-containing transcripts (Moldován et al., 2018a; Tombácz et al., 2019). In this work, we identified 152 5’-truncated transcripts with in-frame ORFs (data not shown).

### Long non-coding RNAs

#### Antisense ncRNAs (asRNAs)

This study detected ten asRNAs each with distinct promoters. bICP22-AS1 and bICP22-AS4 are co-terminal with each-other and overlap the 5’-UTR region of bICP22 (**Figure 1A**). UL28-AS1, and UL47-AS1 overlap the ORFs of the UL28 and the UL47 genes, respectively (**Figure 1B and 1C**). Additionally, a high variety of asRNAs are transcribed from the US region, resulting from premature termination of longer transcripts. US2-AS1, US2-AS2 and US1.67-AS2 are starting at the same TSS and are alternatively truncating isoforms of the very long US2-3-4-C1 complex transcript. In the same way, US2-AS3 is a 3’-truncated isoform of US2-3-4-C2, whereas US2-AS8 is a 3’-truncated isoform of US3-4-L1 (**Figure 1D**). US6-AS3 overlaps three ORFs in antisense orientation (**Figure 1E**). An 1,128 nt-long ORF is located on this transcript but its 376 amino acid long putative protein product does not show similarity to any other proteins in the non-redundant protein databases of NCBI.

**Figure 1.**
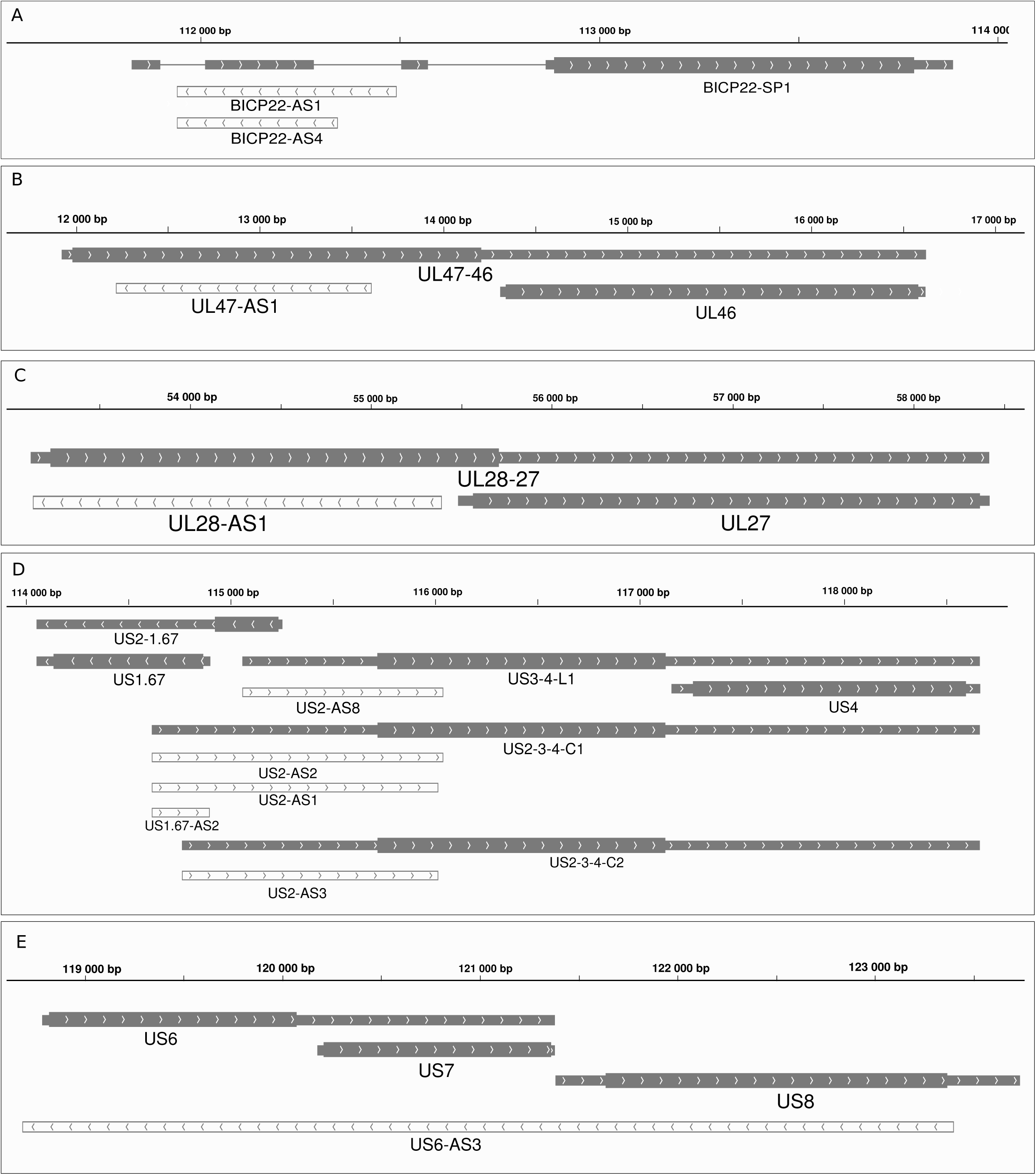
Antisense transcripts of the BoHV-1. Gray rectangles represent mRNAs. Wider overlapping gray rectangles are ORFs. Lines between rectangles represent introns. The orientation of transcripts is represented by arrow heads. White rectangles with gray borders represent antisense transcripts. (A) Two co-terminal antisense transcripts were detected overlapping the 5’ UTR of bICP22-SP1. (B) An antisense transcript overlaps the ORF of ul47. (C) One antisense transcript was detected overlapping the ul28 ORF. (D) Several antisense transcripts were detected overlapping us1.67 and us2. These start at the same TSS as the transcript isoforms of the adjacently located genes us3 and us4. (E) A very long antisense transcript overlaps the us6, us7 and us8 genes.

#### 3’-truncated ncRNAs

***(3tncRNA)*** are the result of premature transcription termination, that lead to the lack of stop codons on the mRNA. We annotated 3’-truncated lncRNAs in 13 genes, totaling 44 transcript isoforms. These RNA molecules lack canonical polyadenylation signal (PAS) and the sequence directly upstream of their termination is GCA-rich, which is not the case for the main transcripts (**Figure 2A)**. The bICP4 gene encodes five 3’-truncated RNAs (**Figure 2B)**, four of which terminate in GA-rich regions. We hypothesize that these loci may act as transcriptional pause sites, that promote termination, cleavage and polyadenylation (Gromak et al., 2006). The UL27 gene produces the highest variation of 3tncRNAs with seven truncated isoforms. Additionally, four of the six 3tncRNAs transcribed from bICP22 are spliced using the same splice sites as the main gene.

**Figure 2.**
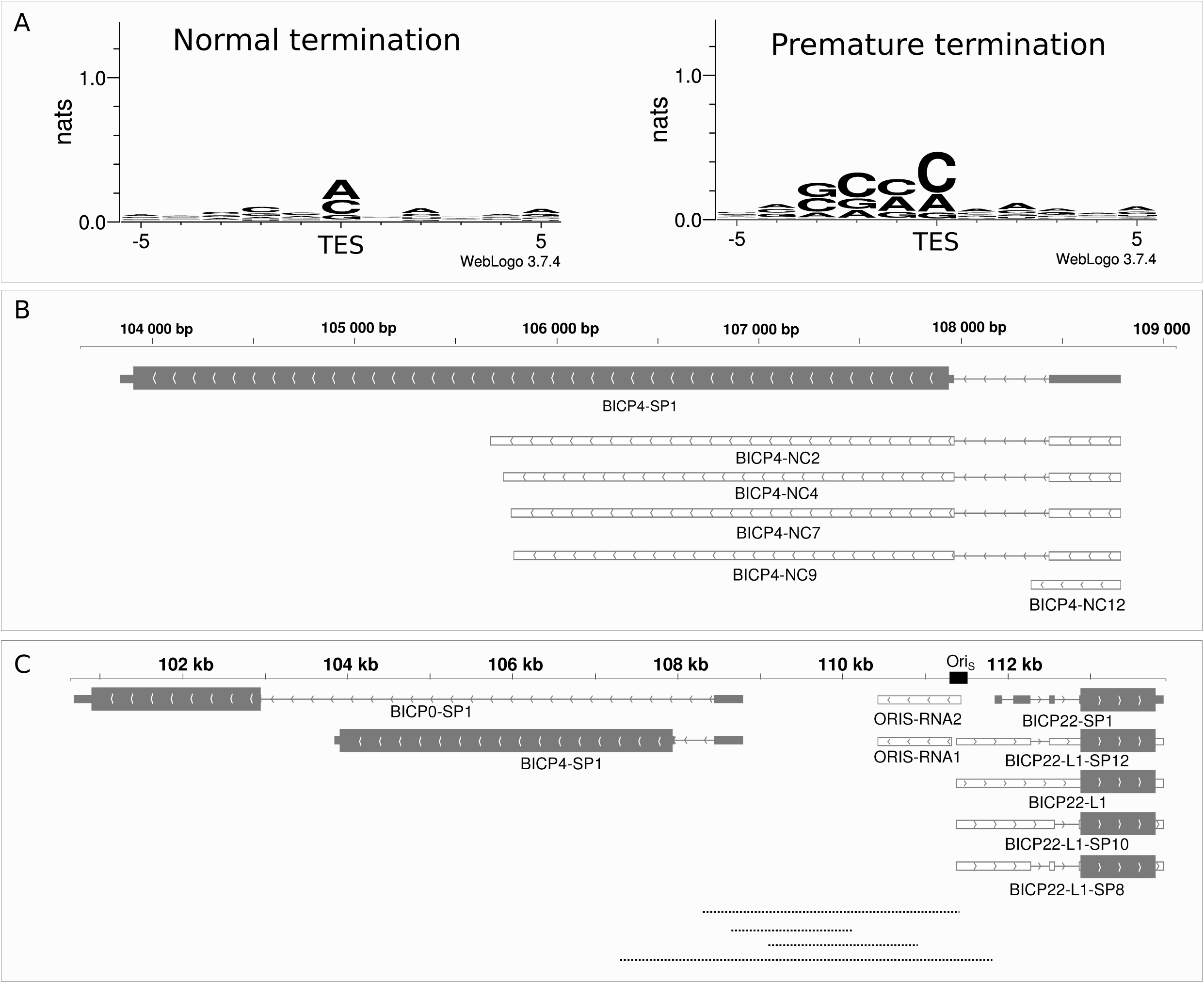
Non-coding transcripts and raRNAs of BoHV-1. (A) The ±5 bp sequence surroundings of the most frequent TESs and the TESs of 3’ truncated transcripts. TESs of prematurely terminating transcripts are characterized by GC-richness immediately upstream of the cleavage site. (B) Gray rectangles represent coding transcripts. The wider overlapping gray rectangle is the ORF. Lines between the rectangles represent introns. The orientation of transcripts is represented by arrow heads. White rectangles with gray borders represent 3’ truncated transcripts. Five 3’ truncated transcripts were detected overlapping the bICP4 gene. (C) Grey rectangles represent transcripts located close to the Ori region. The wider overlapping gray rectangles are ORFs. Lines between the rectangles represent introns. The orientation of the transcript is represented by arrow heads. White rectangles with gray borders represent transcripts overlapping the Ori_S_. Two non-coding RNAs and four 5’-UTR isoforms of bICP22 were detected overlapping the Ori. Dotted lines represent individual reads present in the intergenic region of bICP4-bICP22.

### Replication-associated transcripts

Replication of BoHV-1 starts between 2 to 4h p.i. (Meurens et al., 2004). The viral genome contains a replication origin (OriS) in both repeat regions, but lacks OriL, which is present in its close relative pseudorabies virus (PRV). Replication-associated (ra)RNAs overlap the OriS sequences. We identified six raRNAs: ORIS-RNA1 and ORIS-RNA2, and four 5’-UTR isoforms of bICP22 (bICP22-L1, bICP22-L1-SP10, bICP22-L1-SP12, bICP22-L1-SP8) (**Figure 2C**). The raRNAs were present in low abundance and only at late time points of infection (8h and 12h).

### Transcript isoforms

#### TSS variants and promoters

In this work, we detected 823 TSSs using the criterion for sequencing reads to be present in at least three independent samples. Promoter analysis disclosed 91 TATA consensus sequences at an average distance of 30.6 nt (σ =2.09) upstream from TSS positions and fifteen CAAT consensus sequences at an average distance of 105.8 nt (σ=17.53) **(Table S3)**. The eukaryotic initiator sequence region is poorly conserved in this virus (Py A N U/A) (Javahery et al., 1994). Our analysis revealed a similar consensus but with an enrichment in Gs in the +1 position for TSSs with a TATA-box. TSSs lacking a TATA-box showed an even higher enrichment of Gs in the +1 position followed by a predominance of Gs in the +2 position and an ambiguous +3 position **(Figure 3A)**. This G enrichment has also been described in the initiator element (INR) of HSV-1’s VP5 promoter (Huang et al., 1996; Tombácz et al., 2019). TSSs with a canonical promoter sequence are more abundant at every time point in the infection, than those lacking it **(Figure 3B)**. We identified a putative TSS for RNA1.5 overlapping the genomic junction (Fraefel et al., 1993), which was located at 129,272 nts, and it was found to be co-terminal with the more abundant CIRC RNA. We also detected splice junctions of this transcript located at positions 134,896 and 473. RNA1.5 appears to be a low-abundant CIRC variant with a longer 5’-UTR and an intron.

**Figure 3.**
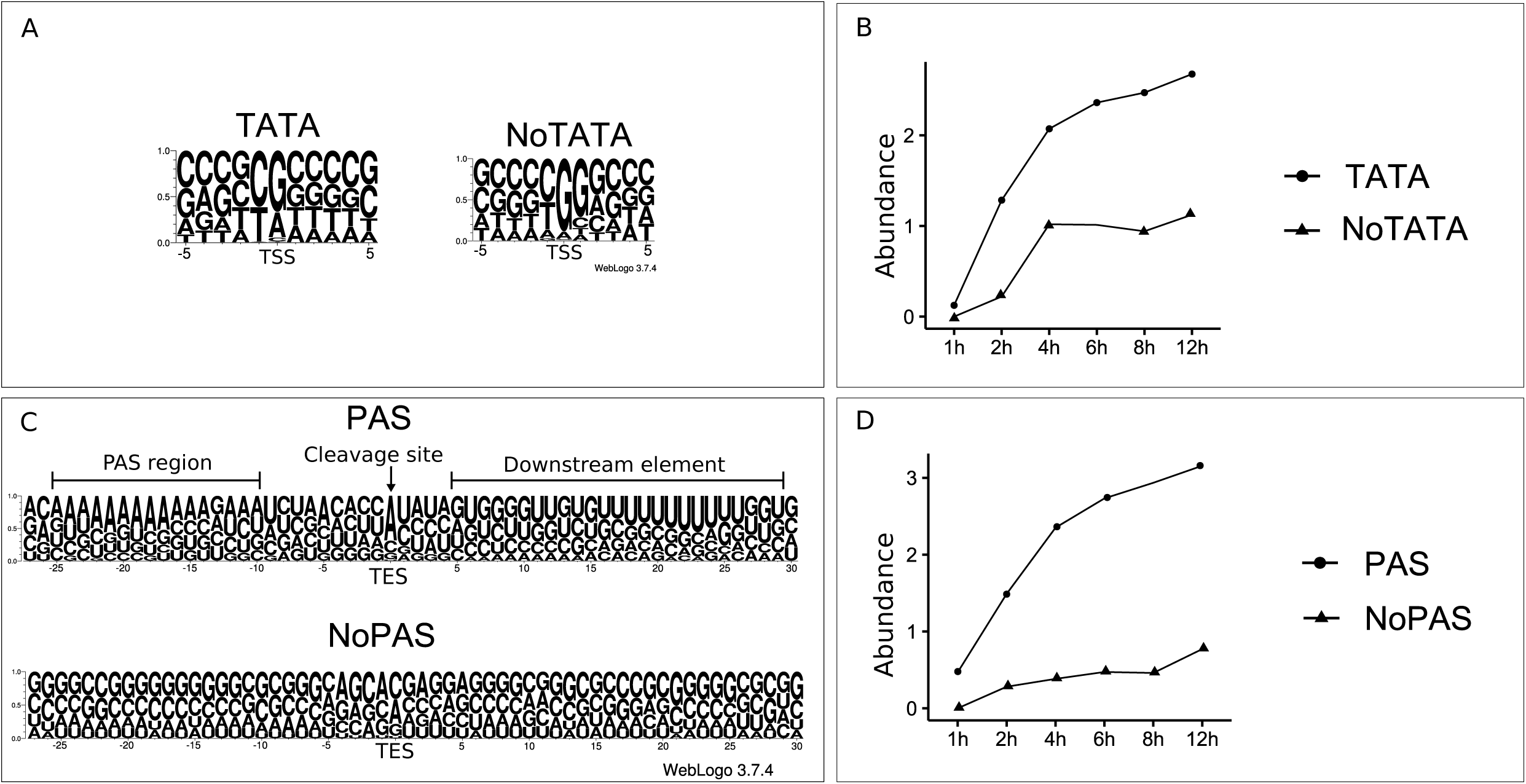
Differential usage of promoters, initiator sites and cleavage sites of the virus. (A) The probability of nucleotides within a ±5 bp interval of the TSSs with or without TATA box promoters. TSSs with an upstream TATA box show a high probability of G/A being located in the TSS position, while the nucleotide in the −1 position being either C or T. TSSs without a TATA box on the other hand are G-rich in the TSS position and the following (+1) nucleotide. (B) The cumulative log_10_ abundance of TSSs with or without a TATA box during the infection. The abundance of TSSs with and without a TATA box increase in the same manner in the IE and E phase of the infection. Following the 4h p.i. time point, the abundance of TATA-less TSSs remains constant whereas the abundance of those with a TATA box is increasing. (C) The probability of nucleotides occurring in the vicinity of the TESs of transcripts with or without a PAS. TESs with a PAS have a canonical cleavage and downstream element (DE) sequence, while TESs without a PAS lack both the consensus cleavage and DE sequence motifs. (D) The cumulative log_10_ abundance of TESs with or without a PAS during the infection. The abundance of TESs with a PAS increases during the entire period of the infection, while the abundance of TESs lacking a PAS shows a steadier increase, with a pronounced steep rise between 8-12 h p.i.

#### TES variants

RNA termination is more precisely regulated than the initiation. This results in a lower TES diversity. Using the above criterion for a LoRTIA transcript, we identified 135 TESs. Our analysis revealed that 48 out of the 135 identified TESs have PAS consensus sequences at an average of 24.5 nt (σ=5.98) upstream of their TESs **(Table S3)**. TESs with PASs have canonical sequence surroundings consistent with the eukaryotic RNA cleavage regions with an orthodox A/C cleavage site and a U/G-rich downstream element, whereas TESs without PAS did not show any consensus sequence motifs at any of the key regions **(Figure 3C)** surrounding the cleavage site. TESs with canonical regulatory sequences are more abundant at every time point of the infection, than those of lacking these sequences (**Figure 3D**). We identified 31 transcripts that terminated in alternative loci compared to their most abundant isoform.

#### Spliced transcripts and splice isoforms

Compared to gammaherpesviruses, alphaherpesviruses have much fewer spliced transcripts. This is indeed true for high-abundance splice isoforms. However, as in HSV-1 (Tombácz et al., 2019), we also detected a number of low-abundance spliced RNA molecules in BoHV-1. In this analysis, we identified 25 introns (23 with GT/AG and 2 with GC/AG) by dRNA-Seq. All of these introns were also detected using cDNA-Seq techniques. We identified all of the previously annotated six introns. However, one of the bICP22 introns spanning from 112,284 to 112,865 (Schwyzer et al., 1994) was present in only two samples represented by merely 4 reads. We detected two novel abundant introns in this region, the first with the same donor and the second with the same acceptor positions as the previously annotated splice sites, but with a 67 nt-long exon between the two (**Figure 4A**). We found a total of 23 novel splice variants with every intron being located in the 5’-UTR region of the transcript. We also located a non-spliced version of the bICP22. Most of the splice isoforms of BICP22 are most abundant at 2h p.i. with the exceptions of bICP22-SP3, bICP22-SP12, bICP22-SP14 and bICP22-SP10, which peak at 6h p.i. (**Figure 4B**). A non-spliced version of UL15 was also present in our samples, although in a lower abundance than its previously annotated spliced variant. In addition, we also detected an alternatively spliced UL15 transcript with the previously annotated intron overlapping its splice donor position at 78,040. This novel intron having a length of 3,579 nts is preceded by a short exon with a start codon. The resulting 128-amino-acid-long putative protein showed no similarity with any of the proteins in the NCBI non-redundant protein database (data not shown). Finally, deleted segments found in cDNA that were absent in dRNA-seq data were only considered as *putative introns* if they were present in at least 3 independent samples. We found a total of 28 putative introns, 23 with GT/AG and 5 with GC/AG consensus **(Table S2)**.

**Figure 4.**
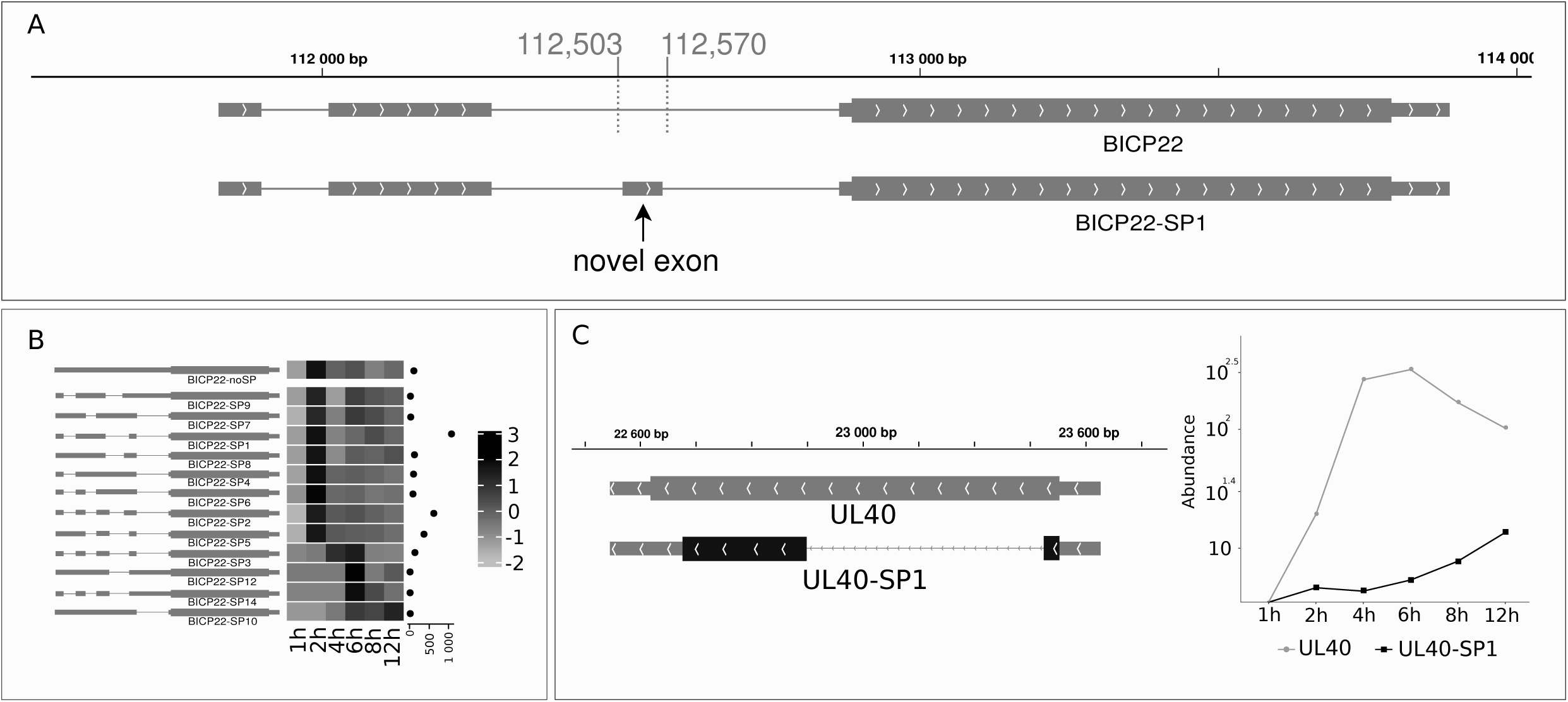
Novel splice variants of BoHV1. Gray rectangles represent coding transcripts. The wider overlapping gray rectangles are ORFs. Lines between the rectangles represent introns. The orientation of the transcript is represented by arrow heads. (A) A novel 67 nt long exon was detected overlapping the second intron of the previously detected bICP22 isoform. The isoform lacking this exon was detected in very low abundance. (B) The change of the expression of specific bICP22 splice isoforms. The scale of the heat map represents Z-scores. The cumulative abundance of each splice isoform is represented by the horizontal dot plot on the right of the heat map. Most of the isoforms show a similar early expression pattern as the most abundant transcript, whereas bICP22-SP3, bICP22-SP10, bICP22-SP12, bICP22-SP14 are expressed more during the later phases of the infection. (C) A novel spliced isoform of UL40 was detected. Splicing causes a frame shift in the *ul40* gene’s ORF (black rectangle), resulting in a 104 aa-long product. The log_10_ abundance of the non-spliced and spliced isoforms is shown on the right. UL40 is characterized by an increased early gene expression and decreasing abundance starting from 6h p.i. Its spliced isoform on the contrary has a low expression in the early phase of the infection, which starts to increase at 4h p.i.

We detected a novel splice variant in a UL40 transcript. The intron spanning from 22,945 to 23,500 produces a frame shift mutation. The predicted 104 aa-long product of the UL40-SP1 showed no significant similarity to any other proteins in the NCBI non-redundant database. This intron is also present in several TSS-variants of the UL40. We observed an increased abundance of the non-spliced isoform until 6 h p.i., followed by a decrease, whereas the abundance of the spliced isoforms started to increase at 4 h p.i. and continued increasing until 12h p.i. (**Figure 4C**).

### Multigenic transcripts

#### Polycistronic transcripts

Herpesvirus genes are organized into tandem gene clusters sharing common transcription termination sequences. It is a general wisdom that in eukaryotes only the most upstream ORFs are translated from polygenic transcripts (Kozak, 1978). In a few examples however, translation from downstream genes of polygenic transcripts has been described in both eukaryotes (Lee, 1991) and their viruses (Ryabova et al., 2002). For example, it has been reported that an upstream ORF (uORF) helps the translation of the downstream ORF36 of Kaposi’s sarcoma-associated herpesvirus (KSHV) by a termination-reinitiation mechanism (Kronstad et al., 2013). Additionally, T3 RNA transcribed from the antisense strand of the KSHV ORF50 gene encodes four micro-peptides, all of which are translated from this polycistronic RNA molecule (Xu and Yuan 2020). We detected a high number of polygenic transcripts using the stringent criteria, including 21 bicistronic, 13 tricistronic, 3 tetracistronic and 4 pentacistronic isoforms (data not shown). To identify the longest transcript isoforms in our samples, we screened the sequencing reads and annotated two more tricistronic and an additional tetracistronic transcripts. These were represented by a single read, thus we deem their start position to be uncertain and hypothesize that they start at the closest TSS. The longest polycistronic transcript annotated by this study is the bicistronic UL37-36 RNA molecule spanning 13,041 nts.

#### Complex transcripts

contain multiple genes of which at least one of the is encoded in the opposite strand. This study revealed a more intricate network of complex (cx)RNA molecules in BoHV-1 than in other herpesviruses (Moldován et al., 2018a; Prazsák et al., 2018; Tombácz et al., 2019), for which the probable reason is technical: in this study we used high-coverage dRNA and dcDNA sequencing techniques. We identified a total of 76 complex transcripts. Twenty-one of these are represented by only one read and were detected by our screening of the longest isoforms. A putative transcript, the bICP22-AT3-SP2 spans almost the entire US region, starting from the TSS of bICP22 and terminating after the ATG of the US9 gene.

### A complex meshwork of transcriptional overlaps

This study revealed an extreme complexity of transcriptional overlaps, which is generated by frequent transcriptional readthroughs and head-to head overlaps produced by divergent genes. Transcriptional readthrough is already a well-known phenomenon between co-terminal tandem gene clusters. Here, we demonstrated that even more distal genes with their own transcription termination are able to carry out occasional transcriptional readthroughs. With a few exceptions (UL30-31, LR-ORF1-bICP0, LR-ORF2-bICP0), canonical transcripts of convergent gene pairs terminate without overlapping each other. Furthermore, we show that every convergent gene produces sporadic readthroughs with a mean overlap size of 153.5 nt (σ=705.6). The most extensive overlap was detected between two putative transcripts UL18-17-16-15-14-13-12-11-C1 and UL15-L4-SP1 with a size of 9,498 nts.

### The effect of BoHV-1 infection on the host cell’s transcriptome

We carried out transcript analysis of the two cell lines on which the virus was propagated. A total of 17,648,405 MinION reads and 14,671 assembled LoopSeq Illumina reads mapped to the *Bos taurus (Bt)* genome, whereas 4,674,656 MinION reads mapped to the *Ovis aries (Oa)* genome. Detailed summary statistics can be found in **Table S1.** We detected a total of 27,290 TSSs, 78,951 TESs and 191,718 introns from *Bt* and 19,222 TSSs 80,542 TESs and 174,392 introns from the *Oa* reads. To eliminate the possibility of false annotation, we filtered TSS, TES and introns that were present in less than two independent *Bt* sample and less than two of the *Oa* samples. As a result, we started transcript annotation with 11,025 TSSs, 21,317 TESs and 139,771 introns for *Bt* (**Table S4**) and 4,939 TSSs 1,562 TESs and 93,406 introns for *Oa* (**Table S5**). LoRTIA produced a total of 227,672 transcripts from the *Bt* and 3,356 transcripts from the *Oa* reads. The median length of *Bt* transcripts was 1,678 nt (σ =2386.465) while the median length of *Oa* transcripts was 1,368nt (σ = 3,810.085).

Seven time points p.i. and a mock-infected sample, all in three replicates, were used to analyze the effect of BoHV-1 infection on the *Bt* cells’ transcript isoforms and gene expression. The replicates yielded highly reproducible data (**Figure S2**).

We detected consensus TATA boxes at a mean distance of 31.148 nt (σ = 2.96) upstream of the *Bt* TSSs. PASs were located at a mean distance of 25.35 nt (σ = 8.26) upstream of the host’s TES. Our data shows no significant alteration in the distance between promoters and TSS and the distance between PASs from TES (**Figure S3A and S3B**). Nor was there any significant modification in the distance between sequence of the ±5 nt surrounding of the TSS and the ±50 nt surrounding of the TES of the *Bt* during the infection (**Figure S3C and S3D**).

Hu and colleagues described alternative splicing and alterations in the polyadenylation of human skin fibroblast cells infected with HSV-1 (Hu et al., 2016). To assess changes in splicing, TSS and TES usage of the host cell during BoHV-1 infection, we evaluated transcripts represented by more than ten reads in the infected samples (n = 69,726) reported by LoRTIA. We detected 130 alternatively spliced transcripts (**Figure S4A**).

FOS, an immediate responder of the stress signaling pathway, is quickly degraded if its third intron is retained (Jurado et al., 2007). We detected a non-spliced variant of FOS in very low abundance and additional splice variants of the transcript lacking the above-mentioned exon, which were present starting from the first hour of the infection (**Figure S4B**). This confirms previous reports on the presence of FOS in the early stages of viral infections (Hu et al., 2016; Rubio and Martin-Clemente, 1999).

The 3’-UTRs of genes often contain miRNA targets, contributing to mRNA degradation. Thus, shorter 3’-UTR length can lead to increased transcript stability (Mayr and Bartel, 2009), whereas longer 3’-UTRs can be targeted by several miRNAs and other trans-acting elements resulting in distinct regulation patterns (Pereira et al., 2017). We detected 72 transcripts with TES located further downstream and 122 with TESs located more upstream compared to transcripts in mock samples.

SOD1 confers protection against oxidative damage (Miao and St. Clair, 2009), including that induced by the IFN-I signaling (Bhattacharya et al., 2015). Two 3’UTR isoforms of SOD1 were detected in infected cells which were shorter than those found in the mock sample (**Figure S4C).**

A previous work described the disruption of transcript termination in the host caused by HSV-1 infection, resulting in extensive overlaps between adjacent transcripts (Rutkowski et al., 2015). The length of polyadenylated transcripts is constant during the infection (**Figure S5**), thus RNAs with disrupted termination are supposedly not polyadenylated. Therefore, we inspected our random primed dataset for reads in intergenic regions. Despite this library yielding a comparable measure of reads mapping to *Bt* (n=2,222,987), we were unable to detect any substantial amount of fragments mapping to intergenic regions.

Using LRS, we were able to differentiate between TSS variants. We detected 80 transcripts with upstream and 142 with downstream TSSs. The 5’-UTR region is important for translation regulation via secondary structures, upstream ATGs and uORFs. Downstream TSSs can also result in truncated ORFs possibly coding for truncated protein products.

### The impact of the virus on host cell gene expression

In this part of the study, we investigated the effect of viral infection on the cultured bovine cells using time-course sequencing analysis. We carried our direct cDNA sequencing using three biological replicates in each of the six time-points and the mock-infected sample. We detected the expression of a total of 8,342 host genes with more than ten transcripts in each of the three replicates. Applying the 0.01 false discovery rate (FDR) threshold, we identified 686 genes that exhibited significantly altered expression levels during virus infection. After clustering the genes by their relative expression profiles, we identified six distinct clusters (**Figure 5A and 5B and Table S6)**. The mean expression profile of the gene clusters indicated four groups of genes (clusters 2-5) that were upregulated, one group that had an initial upregulation followed by downregulation (cluster 1) and a single group of genes where expression levels were steadily downregulated during virus infection (cluster 6). We performed over-representation analysis using the 8,342 expressed genes as reference with the PANTHER software tool. We summarized the results of over-representation analysis with an FDR less than 0.05 using GO (Gene Ontology) biological processes and GO molecular functions annotation datasets in **Table S6**. Over-represented genes were categorized into six functional groups according to the GO database **(Figure 5C)** as follows: 296 genes play a role in cellular metabolism, 257 are involved in transcription and RNA decay, 242 in developmental and morphogenetic processes, 187 in immune response and host defense, 161 in translation and protein folding, whereas 61 genes are specifically associated with in viral transcription related processes. Genes of the first cluster had medium expression preceding the infection (which was transiently slightly upregulated at the 1h, and 2h p.i. time points) followed by downregulation at later measurements. Genes in this cluster were over-represented in pathways controlling a wide variety of developmental and morphogenetic processes. Several genes coding for transcription regulatory proteins present in this cluster show diminishing expression throughout the infection. Genes involved in the cytokine regulation of the immune response and inflammatory processes are also affected.

**Figure 5.**
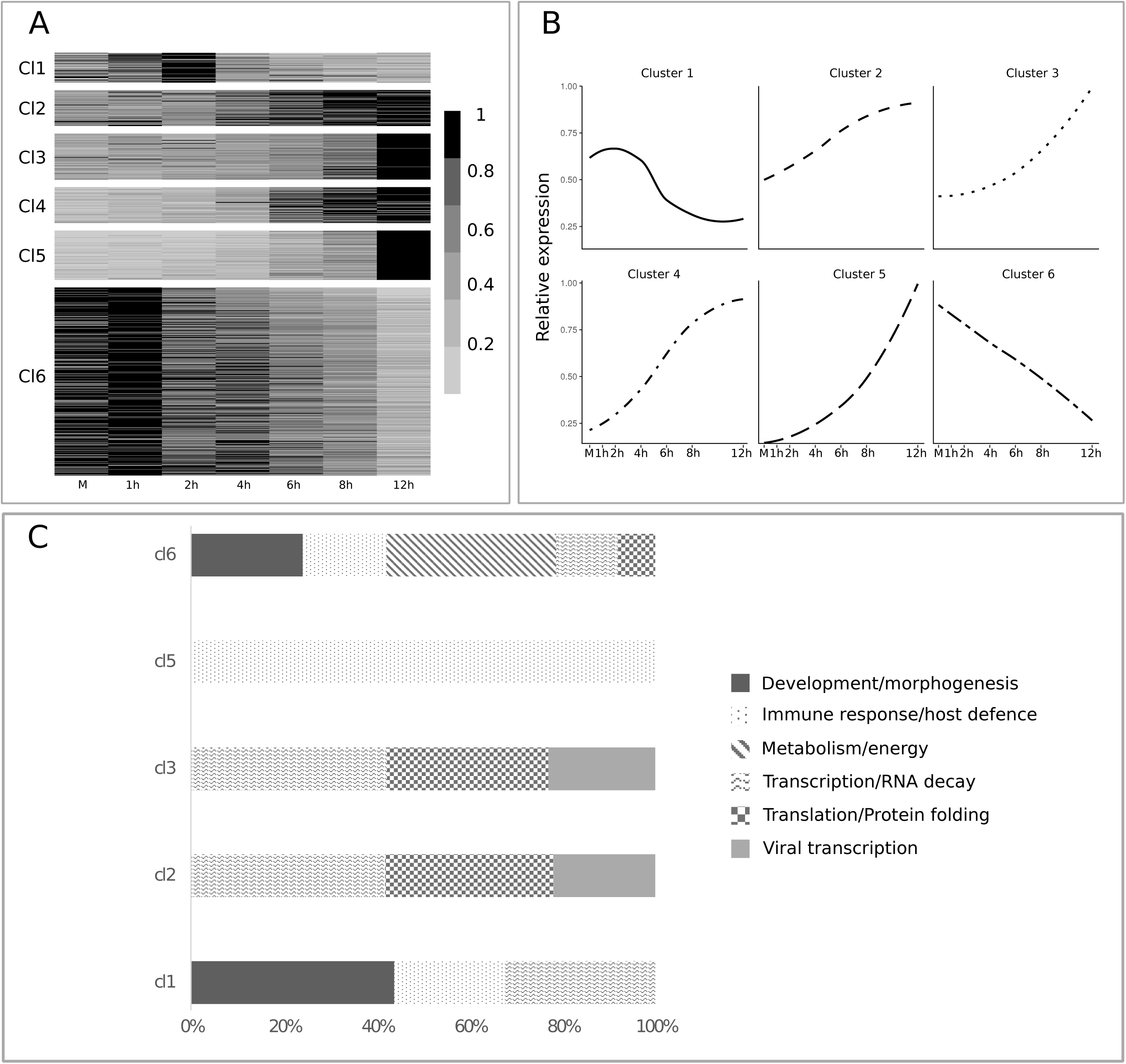
Differential expression analysis of host genes. (A) Relative gene expression represented on a heat map. Six distinct kinetic clusters were identified among host genes. The scale represents the mean relative gene expression. (B) The average change in normalized relative expression of genes present in the six kinetic clusters during infection. (C) The distribution of genes associated with the six functional categories according to gene ontology (GO) between the kinetic clusters.

The second and third cluster of genes are over-represented in functions and molecular processes that can be associated with viral gene expression and the assembly of the virion. An upregulation of genes involved in transcriptional and translational processes, as well as RNA decay was observed. RNA decay can be an immediate response of the host cell to counteract the accumulation of viral transcripts, or it may be an effort of the virus to eliminate competing host mRNAs in order to facilitate the translation of viral transcripts (Moon and Wilusz, 2013; Smiley, 2004). However, some of these GO molecular functions have overlapping sets of genes. For example, the 12 genes (RPS26, RPL5, RPL30, RPS29, RPL31, RPS6, RPL36, RPL37, RPL8, RPS10, RPS21, RPL19) that were significantly upregulated during infection are the members of both the “*viral transcription*” and the “*SRP-dependent co-translational protein targeting to membrane*” pathways. Many of these genes are also members of the “*nuclear-transcribed mRNA catabolic process, nonsense-mediated decay”* pathway. Genes in the fourth cluster were not significantly over-represented in any of the GO molecular functions or GO biological processes. The fifth cluster of genes is over-represented in processes associated with the host defense. Genes present in the type I interferon signaling pathway and other antiviral genes are upregulated during the infection. A huge variety of host genes with high expression preceding viral infection show sharp downregulation during the infection. These are in the sixth cluster and are over-represented in pathways associated with protein folding, the cell cycle regulation and mitochondrial processes including aerobic respiration.

## DISCUSSION

The past decade has witnessed tremendous advances in long-read sequencing. Besides the Pacific Biosciences and Oxford Nanopore Technologies platforms, Loop Genomics has also developed an LRS technique based on single molecules synthetic long-read sequencing. High-throughput LRS techniques are able to read full-length RNA molecules, which led to the discovery that the transcriptomes of examined organisms are much more intricate than previously thought. LRS-based studies have demonstrated that herpesviruses exhibit astoundingly complex transcription profiles (Moldován et al., 2018a; Prazsák et al., 2018; Tombácz et al., 2019). Here we report the assembly and annotation of the BoHV-1 dynamic transcriptome as well as the impact of virus infection on host cell’s gene expression. Our analysis identified a large number of novel transcripts and transcript isoforms. Our results demonstrate that practically every BoHV-1 gene produces transcripts with one or more 5’-truncated in-frame ORFs. It is unknown whether these ORFs are translated, and if so, what their functional significance could be. Several novel splice sites were also identified in this work. This result suggests that splicing in alphaherpesviruses may be more common than previously thought. However, in most cases, splice isoforms are expressed in low abundance relative to the common variant. It can be speculated whether the relative abundance of the splice isoforms is dissimilar in different cell types. We detected a huge variety of bICP22 (homologous to the HSV-1 US1) splice isoforms and found that splicing occurred in the 5’-UTR of the transcript that overlaps two asRNAs. It is possible that these asRNAs contribute to the alternative splicing of bICP22, as suggested in other systems (Krystal et al., 1990; Mihalich et al., 2003). Novel introns in the UL40 and UL15 genes give rise to a change in amino acid composition. The increase of UL40-SP1 following viral replication might play a role in the virus’ transition into post-replication-phase. Replication-associated RNAs have been shown to regulate replication by altering primer synthesis (Tomizawa et al., 1981) or mediate the recruitment of the Origin Recognition Complex (Mohammad et al., 2007). We discovered six replication-associated RNAs (ORIS-RNA1, ORIS-RNA2 and four 5’ UTR isoforms of bICP22), which overlaps the replication origin (OriS) of the virus. ORIS-RNA1 and ORIS-RNA2 are non-coding raRNAs and have no homologous counterparts in closely related viruses. This finding is consistent with observations in closely related herpesviruses.

In HSV-1, a long TSS isoform of ICP4 transcript overlaps the OriS, whereas in PRV the non-coding PTO (Tombácz et al., 2016) and the long TSS variant of the US1 transcript overlap the OriS. It appears to be that raRNAs underwent very rapid evolution, and every species has developed unique solutions for raRNA production. Additionally, two miRNAs have been reported to overlap both raRNAs in an antisense orientation (Glazov et al., 2010), which may play a role in the regulation of raRNAs. Our study demonstrated the existence of very long polycistronic transcripts, and also that genes thought to be present in only bi- or polycistronic RNAs, are also expressed as monocistronic transcripts. A great diversity of transcription initiation is also described in this report. The same ORFs are expressed in a large number of TSS isoforms, with many starting in adjacent genes, and some in more distal genes. A large fraction of this variation is represented by low-abundance transcripts. 5’-UTRs were shown to modulate translation by forming specific structures (Leppek et al., 2018), or through upstream AUGs or uORFs (Calvo et al., 2009; Geballe and Mocarski, 1988; Kronstad et al., 2013). The function of the great extent of TSS variation in BoHV-1 is currently unknown; however, it may lead to post-transcriptional modifications, or differential translation. The prototypic organization of the herpesvirus transcriptome is characterized by the 3’-co-termination of the transcripts produced by tandem genes. Despite this fact, our findings suggest TES variation being substantially lower than TSS diversity. However, transcriptional readthrough is common not only in the parallel, but also in convergent genes. The 3’-UTRs may contain cis-acting elements determining the fate of RNAs after transcription (Matoulkova et al., 2012).

An extremely complex meshwork of transcriptional overlaps was also described. Overlaps are produced by transcriptional read-throughs between tandem and convergent genes or by the mutual utilization of certain genomic loci by divergent genes. Multiple transcriptional read-throughs can result in the generation of complex transcripts. Convergent and divergent transcriptional overlaps produce antisense RNA segments on transcripts. These overlaps may give rise to double-stranded RNAs (dsRNAs), which are targeted by the dsRNA-activated protein kinase R (Dauber and Wolff, 2009) or type I interferon system (Sparrer and Gack, 2015) of the host, thereby effectively reducing viral translation. In a recent publication Dauber and co-workers reported the mechanism of how HSV-1 virion host shut-off (VHS) protein limits the accumulation of viral dsRNAs (Dauber et al., 2019). In light of these findings, it is possible that the frequency of RNAPol-II read-throughs are much higher than expected from the basis of ratio of read-thorough transcripts.

The high-coverage of sequencing reads allowed us to carry out kinetic characterization of viral transcripts. We used direct cDNA data because it produces longer reads than both amplification-based and native RNA sequencing techniques. At the same time, dcDNA-Seq lacks the potential amplification biases causing improper quantitation (Mathieu-Daudé et al., 1996; Polz and Cavanaugh, 1998). LRS allows the distinction between parallel-overlapping RNA molecules and transcript isoforms, therefore we can not only kinetically characterize the genes, but also transcript isoforms.

In this study, we analyzed the effect of viral infection upon host gene expression. We found no significant change in the usage of promoters or PASs of the host genes. However, we observed an altered usage of TESs and splice isoforms. This indicates a modulation of cellular mRNA turnover possibly via miRNA binding sites. The analysis of TSS isoforms suggests that viral infection had an effect on host mRNA translation, potentially through uORFs, or through other cis-acting elements, such as miRNA binding sites of 5’-UTRs (Moretti et al., 2010). Unlike in HSV-1-infected cells (Hu et al., 2016), we found no increase in the rate of transcriptional readthrough events in BoHV-1-infected-cells; albeit our system is not optimal for the investigation of such phenomena.

Based on expression kinetics, we detected six distinct gene clusters that had significantly altered expression. The over representation analysis of these clusters can be summarized in three functional groups. Genes involved in basic cell functions, including morphogenesis, the cell cycle, signaling, catabolic pathways and aerobe respiration, are generally downregulated during viral infection. On the other hand, genes that play a role in viral transcription and translation are upregulated. We also observed upregulation of the genes involved in the cell’s antiviral response. According to our analysis most of these genes could be associated to distinct molecular functions and biological processes indicating general response to virus infection. However, the rest of the unassociated genes could also play key role either in susceptibility or defense to viral infection. Altogether, these analyses and data provide valuable resources for future functional studies, and for understanding how the virus can overcome host defense mechanisms which are vital for the development of novel antiviral therapies.

**Table 1.** Introns annotated in the BoHV1 transcriptome. The read counts are the raw total counts resulting from the LoRTIA analysis.

| Start | End | Length | Read Count | Strand | Junction consensus |
| --- | --- | --- | --- | --- | --- |
| 22945 | 23500 | 555 | 1,941 | - | GT/AG |
| 76461 | 78040 | 1,579 | 44 | - | GT/AG |
| 76461 | 79835 | 3,374 | 2,643 | - | GT/AG |
| 102951 | 108437 | 5,486 | 141 | - | GT/AG |
| 102954 | 108437 | 5,483 | 72 | - | GT/AG |
| 107968 | 108437 | 469 | 427 | - | GT/AG |
| 111899 | 112011 | 112 | 18,853 | + | GT/AG |
| 112077 | 112210 | 133 | 11,660 | + | GT/AG |
| 112077 | 112408 | 331 | 4 | + | GT/AG |
| 112077 | 112443 | 366 | 5 | + | GT/AG |
| 112077 | 112453 | 376 | 9 | + | GT/AG |
| 112095 | 112210 | 115 | 81 | + | GC/AG |
| 112284 | 112502 | 218 | 22,488 | + | GT/AG |
| 112284 | 112524 | 240 | 9 | + | GT/AG |
| 112571 | 112865 | 294 | 24,661 | + | GT/AG |
| 125197 | 125491 | 294 | 23,449 | - | GT/AG |
| 125538 | 125778 | 240 | 11 | - | GT/AG |
| 125560 | 125778 | 218 | 21,948 | - | GT/AG |
| 125619 | 125985 | 366 | 5 | - | GT/AG |
| 125629 | 125985 | 356 | 4 | - | GT/AG |
| 125634 | 125985 | 351 | 6 | - | GT/AG |
| 125852 | 125967 | 115 | 74 | - | GC/AG |
| 125852 | 125985 | 133 | 11,737 | - | GT/AG |
| 126051 | 126163 | 112 | 18,282 | - | GT/AG |
| 129625 | 130094 | 469 | 429 | + | GT/AG |

**Table 2.** Transcript isoform annotated in BoHV-1.

|  |  |
| --- | --- |
| **Non-coding** |  |
| 5' truncated (-TR) | 43 |
| Non-coding | 44 |
| Antisense (-AS) | 10 |
| **Coding** |  |
| Novel ORF | 1 |
| Possibly protein-coding | 152 |
| Long 5'-UTR (-L) | 451 |
| Short 5'-UTR (-S) | 132 |
| Alternative termination (-AT) | 31 |
| Monocistronic | 44 |
| Bicistronic | 21 |
| Tricistronic | 15 |
| Polycistronic | 8 |
| Complex (-C) | 76 |
| Spliced | 19 |
| Non-spliced | 2 |

## DATA AND CODE AVAILABILITY

The sequencing datasets generated during this study are available at the European Nucleotide Archive’s SRA database under the accession PRJEB33511.

The LoRTIA software suite is available on GitHub: https://github.com/zsolt-balazs/LoRTIA.

Our in-house scripts used to generate the descriptive statistics of reads and transcripts, to analyze promoters and to detect transcript isoforms are available on GitHub: https://github.com/moldovannorbert/seqtools.

## ACKNOWLEDGEMENTS

This study was supported by OTKA K 128247 to ZB and OTKA FK 128252 to DT.

## AUTHOR CONTRIBUTIONS

N.M., G.T., G.G. and M.B. carried out bioinformatic analysis of the viral transcripts. Z.M., and T.K. performed bioinformatic analysis of the host transcripts and of the host-virus interactions. Z.C. and D.T. prepared ONT MinION libraries, carried out ONT sequencing and participated in the analysis. D.T. generated Loop-seq libraries and carried out Illumina sequencing. N.M. and G.T. participated in sequencing. Z.C. and V.J. isolated RNA and DNA. M.B. and G.G. maintained cell cultures and participated in nucleic acid isolation and library preparation. V.J., A.H. and Z.Z. propagated the viruses. N.M., Z.M., T.K. and F.M. drafted the manuscript. Z.B. integrated the data, performed the analysis, and wrote the final version of the manuscript. All authors reviewed and approved the final version of the manuscript.

## MATERIALS AND METHODS

### Cells and viruses

Cultured cells were infected with the Cooper isolate (GenBank Accession # JX898220.1) of Bovine Herpesvirus 1.1. Viruses were maintained and propagated in two different laboratories (one in Mississippi State University, Starkville, Mississippi, USA, and the other one in Institute for Veterinary Medical Research, Budapest, Hungary)

#### BoHV-1 from Mississippi (BoHV-1/MS)

Madin Darby Bovine Kidney (MDBK) cells were incubated at 37°C in a humidified incubator with 5% CO_2_, and were cultured with Dulbecco’s modified Eagle’s medium (DMEM) supplemented with 5% (v/v) fetal bovine serum, 100 U/mL penicillin, and 100 µg/mL streptomycin. Cells were either mock-infected or infected with Cooper isolate (GenBank Accession # JX898220.1) of Bovine Herpesvirus 1.1 (BoHV-1) at a multiplicity of infection (MOI) of 5 plaque-forming units (PFU)/cell, incubated at 4°C for one hour for synchronization of infection, and then placed in a 5% CO_2_ incubator at 37°C. Infected cells were collected at 1, 2, 4, 6, 8, and 12 hours post infection (HPI). Each time-point and mock infection consisted of three replicates (n=3). Cells were washed with phosphate buffered saline (PBS), scraped from the culture plate and centrifuged at 300 RPM for 5 minutes at 4°C.

#### BoHV-1 from Hungary (BoHV-1/HU)

The Cooper strain of BoHV-1 was propagated in ovine kidney (OK) cells (ATCC CRL-6551). OK cells were grown to complete confluency in DMEM (Life Technologies) culture medium containing 10% new-borne calf serum (NCS) and antibiotics. Cells were infected with virus stocks at MOI=1, and grown in DMEM without NCS and incubated at 37°C in a humidified 5% CO2-in-air atmosphere incubator. Cells were washed with phosphate buffered saline (PBS), scraped from the culture plate and centrifuged at 300 RPM for 5 minutes at 4°C. Virus stocks were stored at −80°C.

### Virus DNA isolation

#### BoHV-1/MS

The supernatant of cells infected with BoHV-1 at an MOI of 5 for 24 hours was collected and centrifuged at 8,000 RPM for 30 minutes at 4°C to remove cellular debris. The pellet was discarded, and the clarified supernatant was spun at 25,000 RPM in an ultracentrifuge for 2 hours at 4°C with a 30% sucrose cushion to purify viral particles. The supernatant was decanted, and the viral pellet was resuspended in TE buffer (1M Tris-HCl, 0.2M EDTA). Viral particles were lysed with a final concentration of 1% sodium dodecyl sulfate (SDS) at 37°C for 20 minutes before protein digestion with proteinase K (final concentration 0.5 mg/mL) at 60°C for 30 minutes. DNA was then extracted using phenol/chloroform/isoamyl alcohol mixture (25:24:1), pH 8 (Acros Organics, Thermo Fisher Scientific, Fair Lawn, NJ, USA). DNA was precipitated using 3M sodium acetate and 95% ethanol, incubating at −80°C for an hour and was then centrifuged at 14,000 RPM for 30 minutes at 4°C. The pellet was decanted and washed three times with 70% ethanol before one final centrifugation at 14,000 RPM for 15 minutes at 4°C. The supernatant was discarded, and the pellet was suspended in TE buffer.

#### BoHV-1/HU

The supernatant of infected OK cells was centrifuged at 3000 RPM for 5 minutes at 4°C. Virions were separated by ultracentrifugation (Sorwall WX Ultra 90) at 23,500 RPM for 1 hour at 4°C with 30 % sucrose cushion. Viral DNA was extracted using the Qiagen DNA easy Blood and Tissue Kit.

### ONT – amplified genomic DNA sequencing libraries

4 µg viral DNA was fragmented, using Covaris g-TUBE system, at 4200 RPM for 1 minute. 3 µg DNA was used for ONT Genomic DNA by Ligation Kit (SQK-LSK109). DNA strands were repaired by NEBNext FFPE DNA Repair and Ultra II End-prep (New England Biolabs) enzymes. At the end of library preparation, the adapter ligation was performed using NEBNext Quick T4 DNA Ligase, between each steps the library was purified using Agencourt AMPure XP magnetic beads (Beckman Coulter).

### RNA isolation

#### BoHV-1/MS and BoHV-1/HU

RNA from infected and uninfected cells (MDBK cells for BoHV-1/MS or OA cells for BoHV-1/Hu) was extracted using the NucleoSpin RNA kit (Machery-Nagel, Bethlehem, PA, USA), with the lysis step augmented by the addition of proteinase K (final concentration 0.37 mg/mL).

### Poly(A) RNA selection and rRNA depletion

Two selections steps were carried out for total RNAs. The polyadenylated fraction was enriched using Oligotex mRNA Mini Kit (Qiagen), and rRNA depletion was performed using Ribo-Zero Magnetic Kit H/M/R (Epicentre/Illumina).

### ONT - direct RNA sequencing

The BoHV-1/HU samples were used for the preparation of direct RNA (dRNA) libraries using ONT Direct RNA Sequencing Kit (SQK-RNA001). The complementary strand of cDNA was synthetized using SuperScript IV Reverse Transcriptase (Thermo Fisher Scientific) and an RT adapter containing an overhang of 10 Ts, with the addition of 500 ng poly(A)-selected RNA. RT adapters (supplied in the kit) were ligated to the RNA strand using T4 DNA ligase (New England Biolabs). Purification steps were performed using Agencourt AMPure XP magnetic beads (Beckman Coulter) treated with RNaseOUT Recombinant Ribonuclease Inhibitor (Thermo Fisher Scientific). Library concentration was determined using Qubit Fluorometer (v.4.0) and Qubit DNA HS Assay Kit (Thermo Fisher Scientific).

### ONT – direct cDNA sequencing

Non-amplified cDNA libraries were prepared from the mock and six BoHV-1/MS p.i samples in three replicates using the ONT’s Direct cDNA (dcDNA) Sequencing Kit (SQK-DCS109) according to the manufacturer’s instructions. Briefly, first strand synthesis was performed using Maxima H Minus Reverse Transcriptase (Thermo Fisher Scientific) together with SSP and VN primers (supplied in the sequencing kit) and 100 ng of poly(A)-selected RNA for each sample. This was followed by RNA decontamination using RNase Cocktail Enzyme Mix (Thermo Fisher Scientific), and second strand synthesis using LongAmp Taq Master Mix (New England Biolabs). Double stranded cDNA ends were repaired using NEBNext End repair /dA-tailing Module (New England Biolabs) and consecutive sequencing adapter ligation employing the NEB Blunt /TA Ligase Master Mix (New England Biolabs). The enzymatic steps (RT, RNA decontamination and ligation) were carried out in a Veriti Cycler (Applied Biosystems) according to the sequencing protocol. Purification steps were performed using Agencourt AMPure XP magnetic beads (Beckman Coulter) treated with RNaseOUT Recombinant Ribonuclease Inhibitor (Thermo Fisher Scientific). Library concentrations were determined using Qubit Fluorometer (v.4.0) and the Qubit ds(DNA) HS Assay Kit (Life Technologies).

Libraries were barcoded using Native Barcoding (12) Kit (ONT) according to the manufacturer’s instructions.

### ONT – amplified cDNA sequencing libraries

Amplified cDNA libraries were produced for both the BoHV-1/MS and BoHV-1/HU samples using ONT Ligation Sequencing Kit 1D (SQK-LSK109), with the BoHV-1/MS samples being previously pooled in equimolar ratios. Briefly:

#### RT with oligo(dT) primers

50 ng of poly(A)-selected RNA was reverse transcribed using SuperScript IV Reverse Transcriptase (Thermo Fisher Scientific) and oligo(dT) primers (supplied in the kit). The cDNA samples were subjected to PCR using KAPA HiFi DNA Polymerase (Kapa Biosystems) and Ligation Sequencing Kit Primer Mix. End repair, sequencing adapter ligation, library purification and concentration measurements were carried out as described for the direct cDNA sequencing library preparation.

#### RT with random primers

50 ng of ribodepleted RNA was reverse transcribed using SuperScript IV Reverse Transcriptase (Thermo Fisher Scientific) and custom-made primers composed of a random hexamer sequence and one complementary to Ligation Sequencing Kit Primer (supplied in the kit). PCR, end repair, sequencing adapter ligation, library purification and concentration measurements were identical to the oligo(dT) primed RT library. The resulting four libraries were barcoded using the 1D PCR Barcoding (96) Kit (Oxford Nanopore Technologies) following the manufacturer’s instructions.

### LoopSeq single-molecule synthetic long-read sequencing

LoopSeq libraries were prepared from multiplexed 2 h and 12 h post infection samples in three replicates using LoopSeqTM Transcriptome 3x 8-plex Kit. Phasing mRNA protocol was performed according to the manufacturer’s instructions.

### Sequencers

Sequencing of the ONT dRNA, dcDNA and amplified cDNA libraries was performed on R9.4.1 SpotON Flow Cells (ONT). To avoid barcode cross-talk from later time points, mock-infected, 1h and 2h p.i. samples were sequenced separately from other samples. The LoopSeq library was sequenced on an v2 300 flow cell on the Illumina MiSeq system. The two gDNA libraries were sequenced on two R9.4.1 Flongle Flow Cells (Oxford Nanopore Technologies).

### Pre-processing and data analysis

#### Transcriptomic datasets

The MinION data was base called using Guppy base caller v. 3.4.1. with --qscore_filtering. Reads with a Q-score greater than 7 were mapped to the circularized viral genome (NCBI nucleotide accession: JX898220.1) with the Minimap2 software (Li, 2018).

The same reads were also mapped to the host genomes, as follows: the BoHV-1/MS sample was mapped to the genome assembly of *Bos taurus* (GCF_002263795.1) while the BoHV-1/HU sample to the genome of *Ovis aries* (GCA_002742125.1), both using the Minimap2 aligner.

Synthetic long-read data was base called on the MiSeq device using the Real-Time Analysis software with default settings. Base called reads were then processed by the Loop Genomics Pipeline Software with default settings. Synthetic long reads were mapped to the BoHV-1 and *Bos taurus* genomes using Minimap2.

#### Structural analysis

For transcript isoform detection and annotation, mapped reads were analyzed using the LoRTIA software suite v.0.9.9, with the following options:

- For dRNA-seq and dcDNA-seq reads: *-5 TGCCATTAGGCCGGG --five_score 14 -- check_in_soft 15 −3 AAAAAAAAAAAAAAA --three_score 14 -s poisson –f True*.
- For o(dT) and random oligo-primed cDNA reads: *-5 TGCCATTAGGCCGGG -- five_score 14 --check_in_soft 15 −3 AAAAAAAAAAAAAAA --three_score 14 -s poisson –f True*.
- For synthetic long-reads: *-5 ACTAATACTCGGGGG --five_score 16 --check_in_soft 15 - 3 AAAAAAAAAAAAAAA --three_score 14 -s poisson –f True*.

The resulting putative TSSs and TESs were considered as existing if they were detected in at least three independent samples. Deleted segments were accepted as introns (1) if they were present in both dRNA-Seq and one of the cDNA-Seq datasets, (2) if they carried one of the canonical splice junction sequences: GT/AG, GC/AG, AT/AC, (3) if they were shorter than 10 kbps, and (4) if they were more abundant than 1‰. We set a relatively low abundance for acceptance because it may vary in different cell types. The accepted TSSs, TESs and introns were then assembled into putative transcripts using the Transcript_Annotator software of the LoRTIA toolkit. Very long unique or low-abundance reads which could not be detected using LoRTIA were annotated manually. These reads were also accepted as putative transcript isoforms if they were longer than any other overlapping RNA molecule. In some cases, the exact TSSs were not annotated. Finally, a read was considered as transcript if it was present in at least three separate samples. Transcript annotation was followed by isoform categorization according to the following principles: the most abundant transcript containing a single ORF was termed canonical monocistronic transcript, whereas isoforms with longer or shorter 5’-UTRs or 3’-UTRs regions than the canonical transcripts were termed TSS or TES isoforms (variants), respectively. Similarly, transcripts with alternative splicing were named splice isoforms. Transcripts with 5’-truncated in-frame ORF were termed as putative mRNAs. Transcripts with multiple non-overlapping ORFs were designated polycistronic, whereas those with ORFs in different orientation were called complex transcripts. Transcripts with no ORFs or ORFs shorter than 30 nts were named non-coding, except if occurred in front of a canonical ORF in a transcript (These small ORFs were termed uORFs).

#### Analysis of viral gene expression

For the characterization of viral gene expression kinetics, we used dcDNA-seq datasets annotated by LoRTIA. First, we filtered out reads that were confirmed by less than 4 other reads in the dcDNA-seq datasets, then we normalized read counts using a modified version of the Median Ration Normalization of the DESeq2 software suite (Wu et al., 2019). The average normalized read counts of the biological replicates was then considered as the read count of a given isoform.

#### Analysis of host cell gene expression

In order to assess the effect of the infection on the host’s gene expression, we excluded MAPQ=0, secondary and supplementary alignments from all downstream analysis. The reads aligned to the host genome were associated to host genes according to the GCF_002263795.1_ARS-UCD1.2_genomic.gff genome coordinates. Only reads matching the exon structure of the host reference genes (using a +/- 5 base pair window for matching exon start and end positions) were counted. We used edgeR_3.24.3 (McCarthy et al., 2012) with R version 3.5.1 for differential expression (DE) analysis, and filtered out host genes with less than ten reads in any of the three biological replicates. Since we had mock, 1h, 2h, 4h, 6h, 8h, 12h measurements, we used the GLM model (robust=True) and the TMM normalization method in the edgeR analysis. In our model, we tested for DE against mock expression for each time points using data from three biological replicates. To detect genes with significantly changed expression levels, we applied a 0.01 false discovery rate (FDR) threshold, with p-values adjusted by the Benjamini & Hochberg procedure.

#### Pathway analysis of the DE genes

Median of normalized pseudo-counts of DE genes were exported from edgeR. We transformed absolute normalized expressions relative to the highest expression point for each gene to assess which of them had similar expression changes during viral infection. Genes were clustered by their relative expression profile using the amap_0.8-16 R package Kmeans function with the Euclidean distance method. Based on the Calinski criteria, our dataset had an optimal number of cluster number of six. Using the identified subset of genes, we performed overrepresentation analysis for each cluster using the number of expressed genes as reference via the PANTHER (version 14.1 using the 2018_04 dataset release) (Mi et al., 2013) software tool. We summarized the results of our over-representation analysis having an FDR less than 0.05 using the Gene Ontology (GO) biological processes and GO molecular functions annotation datasets.

#### Genomic datasets

MinION data was base called via Guppy base caller v. 3.4.1. using -- qscore_filtering. Reads with a Q-score greater than 7 were assembled using the Flye software (Kolmogorov et al., 2019) with the *--nano-raw -g 164k -m 1000 --meta* options. The assemblies were aligned to the viral reference genome (JX898220.1) using Kalign (Lassmann and Sonnhammer, 2005). The ends of the BoHV-1/MS contig were trimmed, as it was assembled in a concatemer probably resulting from the sequencing of circularized genomic versions (Schynts et al., 2003). Disagreements between the reference and the two assemblies were assessed manually using the UGENE software suite (Okonechnikov et al., 2012). For data visualization we used IGV (Thorvaldsdóttir et al., 2013), WebLogo v.3.0 (Crooks et al., 2004), ComplexHeatmap (Gu et al., 2016), pheatmap_1.0.12 and ggplot2 (Wickham, 2016) R packages. For a schematic representation of our workflow see **Figure S1**.

## SUPPLEMENTARY TABLES LEGENDS

**Table S2. TSSs, TESs and introns of BoHV-1 detected by LoRTIA.**

**Table S3. Promoters and polyadenylation signals of viral transcripts.**

**Table S4. TSSs, TESs and introns of the MDBK cell line detected by LoRTIA.**

**Table S5. TSSs, TESs and introns of the OK cell line detected by LoRTIA.**

**Table S6. The relative expression, GO biological processes and molecular functions of the MDBK cell line’s genes.**

