## Supplementary info 1 for "Time-course Profiling of Bovine Herpesvirus Type 1 and Host Cell Transcriptomes using Multiplatform Sequencing"

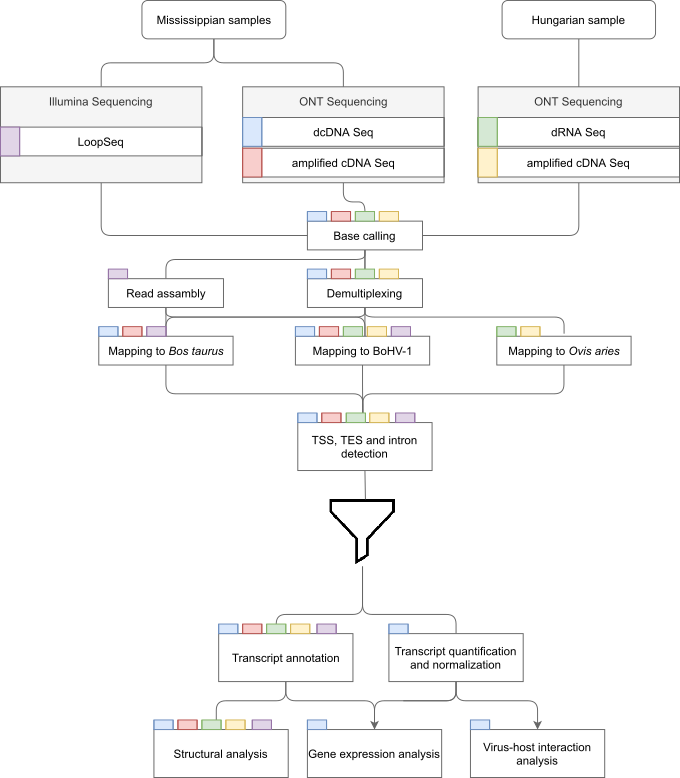

**Figure S1.** **Schematic representation of the workflow.** Colored rectangles represent the sequencing libraries. Steps of data analysis are shown in rectangles, with the libraries undergoing the given step indicated by the tab color. The funnel symbol represents TSS, TES and intron filtering.

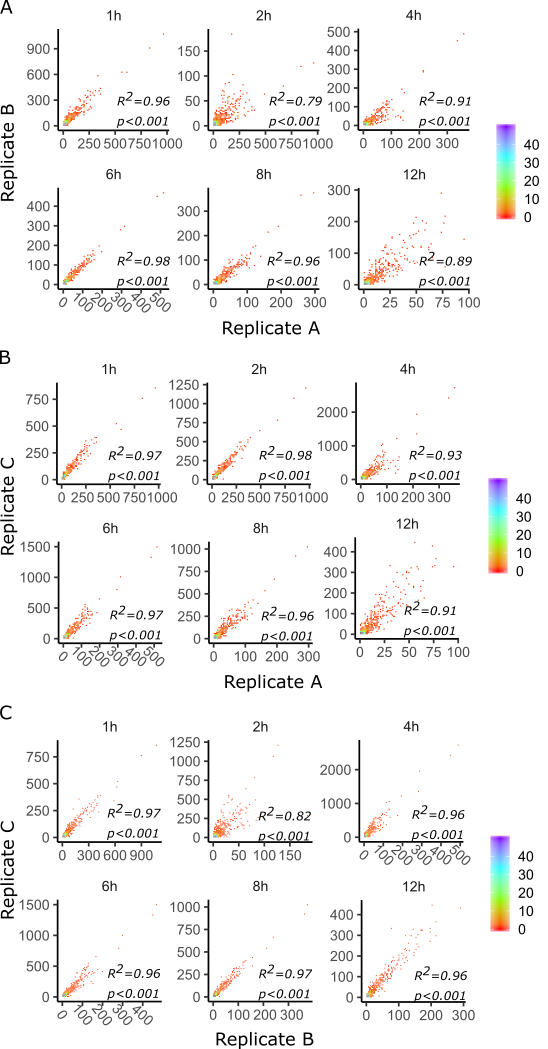

**Figure S2. Reproducibility of the results.** Pearson’s correlation of the read counts for (A) Replicate A and B, (B) Replicate A and C and (C) Replicate B and C. Data density is show in colors on the right of the scatter plots, where blue and purple represents mora abundant data points. A high correlation was detected among the read counts of the replicates.

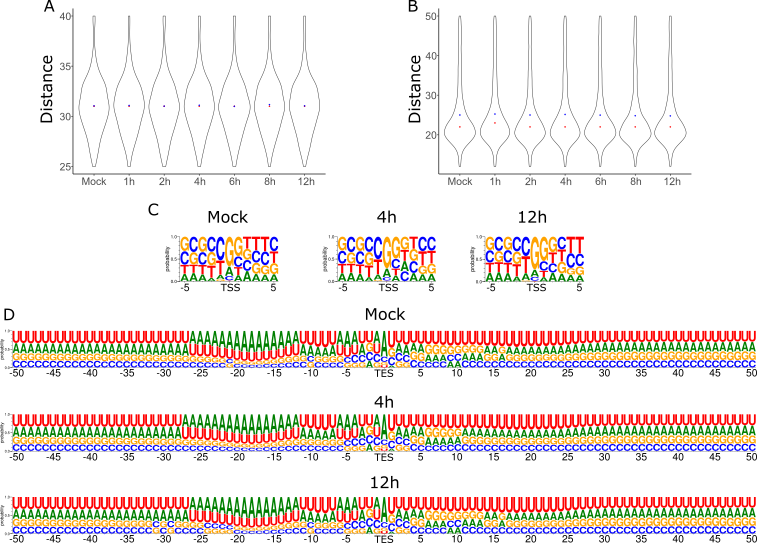

**Figure S3. The effect of the infection on the host’s transcript initiation and termination.** (A) No significant change was detected in the mean (blue dot in the violin plot) nor median (red dot in the violin plot) distance of the TATA boxes from the TSS. The distance is presented in bp. (B) or the mean (blue dot) nor median (red dot) distance of the polyadenylation signal from the TES. The distance is presented in bp. (C) The infection has no observable effect on the initiator sequence (D) nor the surrounding sequence of the polyadenylation and cleavage site as shown by the probability of nucleotides.

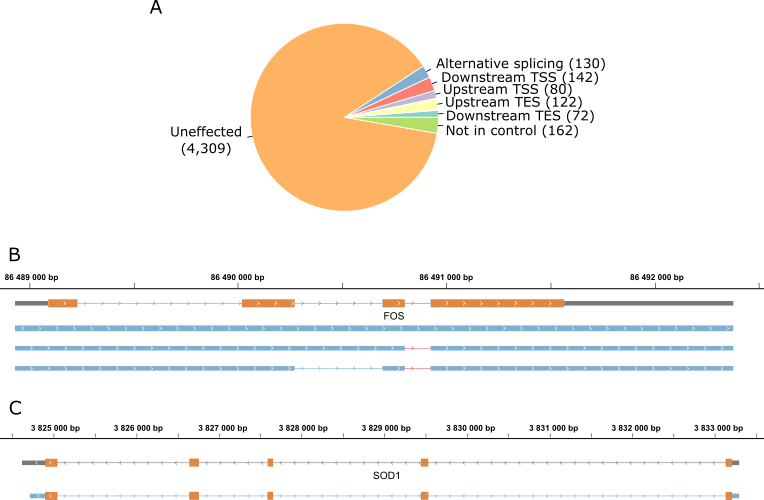

**Figure S4. The effect of the infection on the host’s transcript isoforms.** (A) A total of 546 transcript isoforms were detected with at least ten reads in the p.i. *Bt* datasets compared to the transcripts present in the mock dataset. (B) The blue rectangles represent a novel non-spliced and two novel splice variants of the FOS transcript. Introns are represented by lines between the rectangles. The intron which if retained is responsible for increased degradation of FOS is shown in red. Orange rectangles represent the ORF. (C) The blue rectangles represent the novel TES isoform of SOD1, lines represent introns, while orange rectangles represent the ORF. Isoforms of SOD1 with an upstream TES were detected during the infection but not in the mock samples.

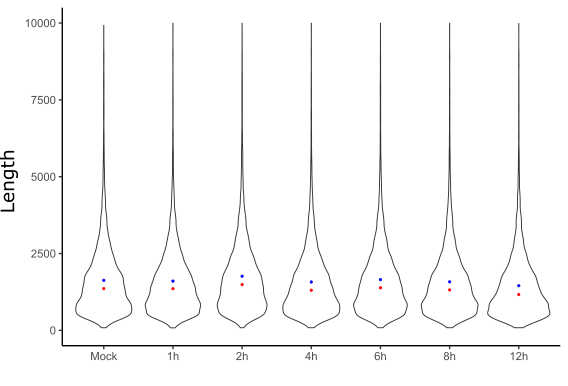

**Figure S5. The length of MDBK cell transcripts isolated from different p.i. time points.** Blue dots represent the mean while red dots the median lengths. No significant change can observed in transcript lengths during the infection. The length is presented in bp.

**Table S1.** **Summary statistics of the BoHV-1 and host sequencing grouped by library types.** For the MinION platform, reads with a quality score of 7 or higher, whereas for the Illumina platform, reads with a Q-score of 30 were selected for analysis. The table contains contig counts for Illumina LoopSeq, which were assembled from a total of 36,627,542 sequencing reads.

| **Library type** | **Raw read count** | **Mapped read count**  **(primary mappings)** | | | **Mean mapped read length**  **(primary mappings)** | | |
| --- | --- | --- | --- | --- | --- | --- | --- |
|  |  | **BoHV-1** | **B. taurus** | **O. aries** | **BoHV-1** | **B. taurus** | **O. aries** |
| ONT dRNA | 916,140 | 516,294 | - | 249,035 | 980.525 | - | 873.931 |
| ONT dcDNA ONT o(dT) | 15,379,697 | 1,830,137 | 13,126,238 | - | 1,152.544 | 1,179.308 | - |
| cDNA ONT o(dT) | 4,717,241 | 688,617 | 2,299,180 | 1,541,723 | 677.384 | 1,456.542 | 1,023.767 |
| cDNA ONT random | 5,650,274 | 352,903 | 2,222,987 | 2,883,898 | 662.14 | 1,104.522 | 791.055 |
| Illumina LoopSeq | 27,098 | 12,618 | 14,671 | - | 642.3 | 853.786 | - |
